# MyGene.info and MyVariant.info: Gene and Variant Annotation Query Services

**DOI:** 10.1101/035667

**Authors:** Jiwen Xin, Adam Mark, Cyrus Afrasiabi, Ginger Tsueng, Moritz Juchler, Nikhil Gopal, Gregory S. Stupp, Timothy E. Putman, Benjamin J. Ainscough, Obi L. Griffith, Ali Torkamani, Patricia L. Whetzel, Christopher J. Mungall, Sean D. Mooney, Andrew I. Su, Chunlei Wu

**Affiliations:** Department of Molecular and Experimental Medicine, The Scripps Research Institute, 10550 North Torrey Pines Road, La Jolla, CA 92037, USA; Current address: Avera Cancer Institute, 11099 North Torrey Pines Road, La Jolla, CA 92037, USA; Department of Biomedical Informatics and Medical Education, The University of Washington, Box SLU-BIME 358047, Seattle, WA 98195, USA; McDonnell Genome Institute, Washington University School of Medicine, 4444 Forest Park Ave, St. Louis, MO 63108, USA; Department of Integrative Structural and Computational Biology, The Scripps Research Institute, 10550 North Torrey Pines Road, La Jolla, CA 92037, USA; The Scripps Translational Science Institute, The Scripps Research Institute, 10550 North Torrey Pines Road, La Jolla, CA 92037, USA; Center for Research in Biological Systems, University of California San Diego, 9500 Gilman Drive, La Jolla, CA 92093, USA; Lawrence Berkeley National Laboratory, 1 Cyclotron Road, Berkeley, CA 94720, USA

## Abstract

MyGene.info and MyVariant.info provide high-performance data APIs for querying gene and variant annotation information. They demonstrate a new model for organizing biological annotation information by utilizing a cloud-based scalable infrastructure. MyGene.info and MyVariant.info can be accessed at http://mygene.info and http://myvariant.info.

## Content

The accumulation of biomedical knowledge is growing exponentially. There have been tremendous efforts seeking to structure research findings as annotations on various biological entities (e.g., genes, genetic variants, pathways). However, these annotations are fragmented among many resources that range greatly in terms of size, funding, and visibility (e.g., Ensembl^1^, Uniprot^2^, PROSITE^3^ and Reactome^4^). Tools for knowledge integration enable more efficient analysis of genome-scale data sets and discovery of relationships between biological entities.

Bioinformaticians facing data integration problems generally pursue one of two strategies: data warehousing and data federation. Data warehousing involves downloading flat-files from various sources, writing parsers to process the files, and then loading the parsed data in a local database. This strategy has the advantage of very high performance, but also requires significant initial effort to write the parsers and ongoing effort to keep the resource up to date. On the other hand, data federation works by accessing remote data resources through web services. Federated data solutions are always up to date, but extra care is required to maintain the links, and large queries may take a long time to return due to server and network limitations.

Here we present an alternative solution for integrating annotations on genes and human variants. MyGene.info and MyVariant.info are open source, high-performance, and continuously-updated data application programming interfaces (APIs) for accessing comprehensive reports of gene and variant annotation. These resources are offered as cloud-based web service endpoints with the goal of providing “Annotation as a Service”. MyGene.info and MyVariant.info are centralized repositories to store dispersed annotation data.

Other centralized resources for gene and variant annotations currently exist for genes (e.g.,Bioconductor AnnotationData Packages^5^ and Biomart^6^) and variants (e.g., ANNOVAR^7^). Relative to these existing tools, MyGene.info and MyVariant.info have several advantages. First, a local database is not required, reducing set-up, administration and maintenance costs. Second, we provide a high-performance API that allows real-time queries in analysis pipelines or web applications.

MyGene.info is maintained as a comprehensive and up-to-date repository for gene annotations. It integrates data from large, centralized databases as well as small, specialized sources. Each data source has its own data importer, converting external data sources to a list of objects in JSON format. Each individual JSON object uses the NCBI’s gene ID^8^ as the preferred primary key. The output of each parser is stored in a MongoDB instance with a timestamp recorded for each individual annotation object, and all objects with the same primary key are combined together into a single annotation object. In addition, we have built a scheduling system that automates the updates for each data source according to its own schedule. Currently, MyGene.info provides over 200 gene-specific annotation fields, covering more than 13 million genes for 15,000 species (http://mygene.info/metadata).

MyVariant.info is built with a similar design and architecture as MyGene.info, while focusing specifically on annotations of human genetic variants. We utilize the nomenclature from the Human Genome Variation Society (HGVS) to define the primary key in MyVariant.info (see methods for specific rules). To prevent incorrect usage of HGVS IDs that could lead to potential errors in clinical interpretation, we also developed and implemented a variant validation function to ensure that all variant IDs included in MyVariant.info strictly follow HGVS guidelines. Currently, MyVariant.info contains over 500 variant-specific annotation types from dozens of resources, covering more than 316 million unique variants (http://myvariant.info/metadata).

The performance, scalability and stability are three key features of a successful web-service provider. We built an Elasticsearch-based cluster to index the underlying JSON objects for both MyGene.info and MyVariant.info. This indexing engine provides both superior query performance and rich query syntax to handle a large amount of concurrent queries for a variety of use cases. The Elasticsearch cluster also comes with inherent scalability, so that we can dynamically adjust the size of the cluster to accommodate increased bandwidth as needed. For example, The MyGene.info system is currently hosted on the Amazon EC2 platform with four moderate servers, which based on our tests, can handle traffic from >5000 concurrent users for approximately 10,000 requests per minute. Greater than 95% of actual user requests take less than 30ms to process (see Supplementary Fig. 1). This dynamic cluster setup also promotes stability of these services by allowing us to perform maintenance on individual nodes without bringing down the whole system.

Since the release of the v2 API (July 2013), MyGene.info has accumulated over 120 million requests, and currently serves more than 2 million requests per month. Primary users include public resources like BioGPS^10^, Monarch Initiative^11^ and CIViC (https://civic.genome.wustl.edu/). In addition, numerous individual users incorporate MyGene.info into their bioinformatics analysis pipeline. According to our usage monitoring, approximately 40% of traffic comes from our BioGPS application, while 60% of traffic comes externally from over 5000 unique IP addresses. The MyVariant.info API was launched in June 2015. To date, MyVariant.info has accumulated over 790,000 user queries.

To demonstrate their utility, we used MyVariant.info and MyGene.info to re-implement a typical analysis pipeline for interpreting exome sequencing results and identifying candidate genes for a rare Mendelian disease. In 2010, Ng. et al identified *DHODH* mutations as the genetic cause for Miller Syndrome^12^. In their exome analysis, genes with nonsynonymous (NS) variants, splice acceptor and donor site mutations (SS), and coding indels (I) were first identified. Next, they filtered for genes containing NS/SS/I variants in all four sequenced samples. Previously observed variants in dbSNP129^13^, the 1000 Genomes Project^14^ or Hapmap were excluded. PolyPhen predictions^15^ were used to prioritize variants that were predicted to be damaging. This process undoubtedly involved downloading, parsing and analyzing annotation data from multiple databases, representing a significant investment of time and effort.

Using the MyGene.info and MyVariant.info R packages alone, we are able to implement an updated version of this pipeline in less than 100 lines of code, requiring no local installation of variant annotation databases or software tools (https://github.com/sulab/myvariant.info/blob/master/docs/ipynb/myvariant_R_miller.ipynb, see Fig. 1 and Supplementary Note 1). We first filtered for NS/SS/I variants and removed variants observed in the 1000 Genomes project (as in Ng et al.). We also incorporated an allele frequency filter based on data from the Exome Aggregation Consortium (ExAC, http://exac.broadinstitute.org), filtered for candidate genes involved in metabolic processes (“GO:0008152”) based on Gene Ontology annotations, and ranked candidate genes based on Combined Annotation Dependent Depletion (CADD) score^16^ (an estimate of pathogenicity). After implementing this workflow for Miller Syndrome study, we were left with only 5 candidate genes, including the causal gene *DHODH*. In addition, since MyVariant.info contains comprehensive and up-to-date variant annotations, it offers users the flexibility to further tailor this workflow based on other annotation fields (e.g., sift score^17^, polyphen score^15^, clinical significance from ClinVar^18^, etc.).

**Figure 1.**
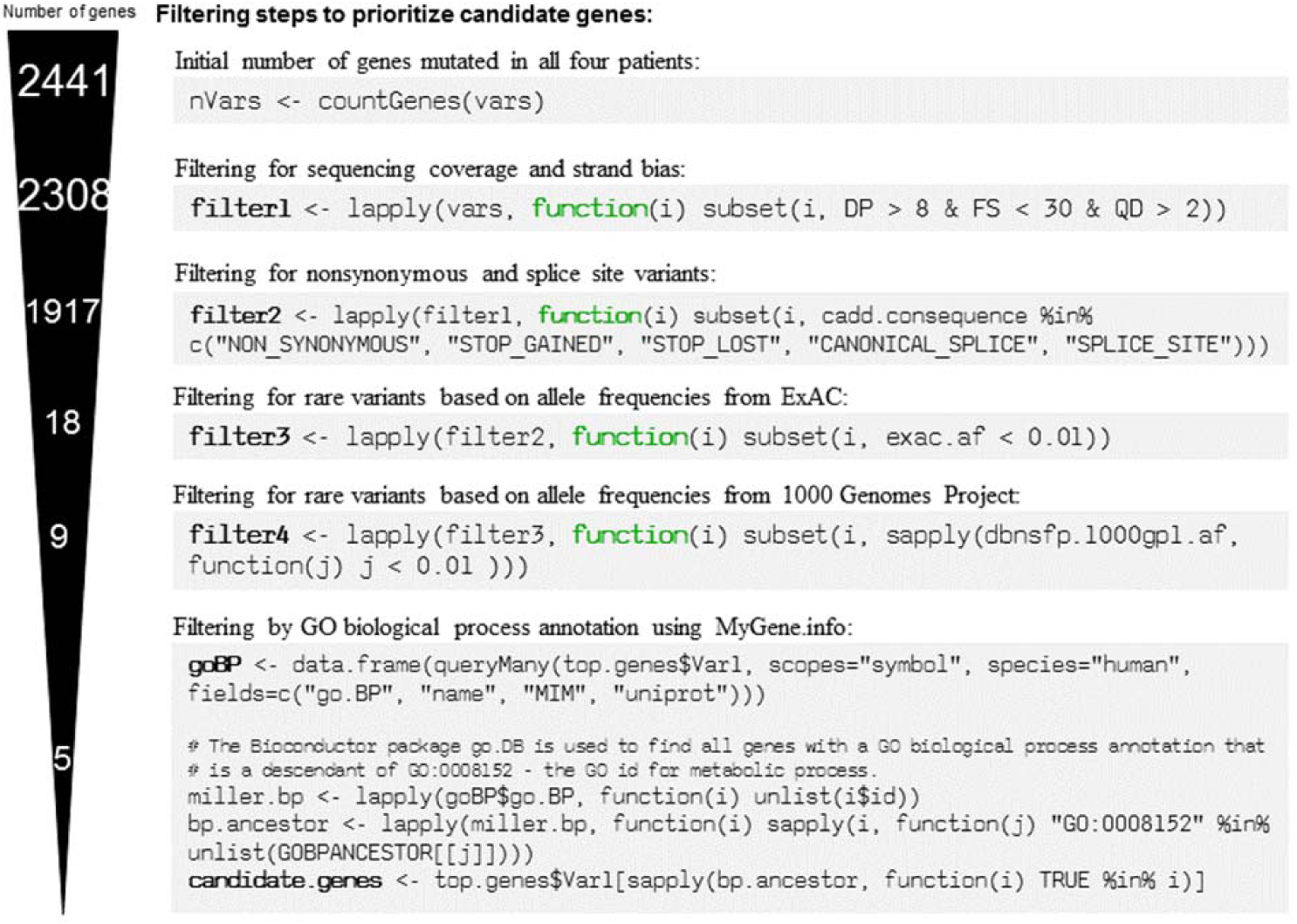
Five filtering steps are shown for the re-implemented workflow to prioritize candidate genes from Miller Syndrome study^12^ using MyVariant.info and MyGene.info web services. Selected R code are displayed for each filter step, using *myvariant* and *mygene* Bioconductor packages. And the number of candidate genes left at each filtering step is displayed at the left side. The full code are available at https://github.com/sulab/myvariant.info/blob/master/docs/ipynb/myvariant_R_miller.ipynb, and also in Supplementary Note 1.

The utility of MyGene.info and MyVariant.info extend beyond this particular pipeline for exome sequencing analysis. Users can search for genes or variants using a wide variety of identifiers, and annotations can be retrieved for either single entities or lists. Users can also perform data-dependent queries (e.g., to find all variants with ExAC allele frequencies below 0.05 in the gene *BRCA1)*. Queries can be performed through the web-based API, or using data access libraries for R or python. These tools are flexible enough to incorporate into custom workflows, as well as responsive enough to perform real-time queries from other web applications. Finally, although MyGene.info and MyVariant.info focus on genes and genetic variants, the underlying open-source infrastructure is extensible to any type of biological entity.

## Methods

### Data Sources

Currently, MyGene.info contains data for ~200 annotation fields that were retrieved from 8 public databases, and MyVariant.info contains data for ~500 annotation fields that were retrieved from 14 public databases (Supplementary Table 1). Our scheduler checks every data resource on a weekly basis, detects any changes to the source files, and applies incremental updates to our live servers. Annotation data from different databases exist in different formats, e.g., VCF, XML and TSV. We wrote an individual data parser for all annotation data sources. Data parsers automatically import data from raw sources to facilitate regular updates where possible. All parser code for MyGene.info is available at https://bitbucket.org/sulab/mygene.hub/src/default/src/dataload/sources/. All parser code for MyVariant.info is available at https://github.com/sulab/myvariant.info/tree/master/src/dataload/contrib.

### Data Integration

The output of each data parser is a list of JSON objects. Each object contains a ‘_id’ field as the primary key, which uniquely identifies a biological entity. MyGene.info uses NCBI gene ID^8^ as the preferred primary key, although Ensembl gene ID^1^ is used when no mapping to NCBI gene is available.

For primary keys for variants, we used the nomenclature defined by the Human Genome Variation Society (HGVS)^9^, as it is a recommended and widely-accepted standard for describing variants. HGVS nomenclature allows multiple names to describe the same variant based on different reference sequences (e.g. genome assembly, transcript or protein sequences). To define unique primary keys, we only use the HGVS names based on the most commonly used reference genome assembly (currently hg19), and use *chr1, chr2,…, chr22, chrX, chrY* and *chrMT* to represent chromosomes (e.g., chr11:g.111959693G>T, more examples in Supplementary Table 2). Although the primary keys of variant objects are based on genomic reference sequences, other valid HGVS names corresponding to alternate reference sequences (e.g., NC_000011.9:g.111959693G>T, NM_003002.2:c.274G>T) are also stored in each variant object and are indexed for queries.

We implemented a scheduling system to automate the updates for each data source. Currently, both MyGene.info and MyVariant.info are updated weekly. The output of each parser is stored in a MongoDB instance with a timestamp recorded for each individual annotation object, and all objects with the same primary key (‘_id’ field) are combined together into a single annotation object (see examples at Supplementary Fig. 2). This setup is advantageous as it ensures the independent processing of each annotation source. Any single failure in the update process will not break the entire merging process, as in the case when a source file format changes and breaks a data parser. In that case, the last successful version of that failed source will be used until the parser is adapted to the changes.

After the merging process, each JSON object contains all variant annotations aggregated from multiple sources. We then use Elasticsearch to index all fields within an annotation object, so that users can make queries to retrieve annotations for their relevant genes or variants. Elasticsearch is a highly scalable, open-source, full-text search and analytics engine. It provides a rich query syntax and inherent scalability to handle large-scale data queries in real time.

### Application Programming Interfaces

On top of Elasticsearch, we built REST-based web services using the Tornado web framework. Tornado is a Python-based web framework built upon asynchronous networking technology that can provide tens of thousands of concurrent connections with a moderate server.

MyGene.info provides two simple-to-use REST-based web services: a gene query service and a gene annotation service. The gene query service allows users to query for gene annotations using any identifier or keyword, while the gene annotation service provides a convenient way to retrieve gene-centric annotations when gene IDs (NCBI gene IDs or Ensembl gene IDs) are available.

MyVariant.info also provides two REST-based web services: a variant query service that returns matching variant objects based on user queries, and a variant retrieval service that returns the matching variant object(s) for a given ID (HGVS names, rsids, etc).

Batch mode is supported by both services for querying a large list of IDs or query terms in one request. Both query services provide rich query syntax suitable for a variety of use cases. For example, users can query for matching variant annotation objects by various criteria, such as genomic ranges, prediction score cutoffs, exact field matching, keyword search in a text field, etc. Users also have the option of specifying the subset of fields they want to return if they do not require the entire annotation objects. More complicated queries can be constructed by combining multiple query terms with Boolean operators, or conducting faceting, aggregations for special use-cases. Information on the types of queries that are enabled by MyGene.info can be found at http://docs.mygene.info/en/latest/doc/query_service.html. Information on the types of queries that are enabled by MyVariant.info can be found at http://docs.myvariant.info/en/latest/doc/variant_query_service.html.

### Use Case Demonstration

We reanalyzed the exome sequencing data generated by *Ng et al* for their Miller Syndrome study^12^. Genomic DNA from four patients were sequenced in this study. FASTQ files were processed according to the GATK best practices^19^. Individual samples were aligned to the hg19 reference genome using BWA 0.7.10. Variants were called using GATK 3.3 HaplotypeCaller and quality scores were recalibrated using GATK VariantRecalibrator. The Bioconductor packages *myvariant* and *mygene* were utilized to demonstrate a streamlined application for variant filtering and prioritization of candidate genes in rare Mendelian disorders.

### Python and R Clients

A Python client and R client are available for both MyGene.info and MyVariant.info. The Python client for MyGene.info can be downloaded at https://pypi.python.org/pypi/mygene. The R client for MyGene.info is released as part of Bioconductor at http://bioconductor.org/packages/release/bioc/html/mygene.html. The Python client for MyVariant.info can be downloaded at https://pypi.python.org/pypi/myvariant/ and the R client for MyVariant.info is also released as part of Bioconductor at http://bioconductor.org/packages/release/bioc/html/myvariant.html.

### Source Code

MyGene.info and MyVariant.info are both open source projects (Apache licensed). The source code for these projects can be found at https://bitbucket.org/sulab/mygene.info (MyGene.info web frontend), https://bitbucket.org/sulab/mygene.hub (MyGene.info data backend) and https://github.com/sulab/myvariant.info (MyVariant.info).

## Acknowledgements

Exome sequence data from two sibs with Miller Syndrome and two unrelated affected individuals was provided by Ng et al through the database of Genotypes and Phenotypes (dbGaP) under accession phs000244.v1.p1.Authors also acknowledge the valuable input from Dr. Robin Haw.

This work was supported by the US National Institute of Health (grants U01HG008473 to C.W., GM083924 and U54GM114833 to A.I.S., U01HG006476 to A.T., K22CA188163 to O.L.G). This work was also supported by the Scripps Translational Science Institute, an NIH-NCATS Clinical and Translational Science Award (CTSA; 5 UL1 RR025774).

## Author Contributions

A.I.S and C.W. conceived of the project. J.X., A.M., C.A. and C.W. implemented the core software. G.T. coordinated the project out-reach. M.J., N.G., G.S.S., T.E.P., B.J.A., O.L.G., A.T., P.L.W., C.J.M. and S.D.M contributed either code or data in the project. J.X, A.M., A.I.S. and C.W. wrote the manuscript with input from all authors. A.I.S, S.D.M and C.W. directed this project.

